# Oxytocin gene networks in the human brain: A gene expression and large-scale fMRI meta-analysis study

**DOI:** 10.1101/149526

**Authors:** Daniel S. Quintana, Jaroslav Rokicki, Dennis van der Meer, Dag Alnæs, Tobias Kaufmann, Aldo Córdova Palomera, Ingrid Dieset, Ole A. Andreassen, Lars T. Westlye

## Abstract

Oxytocin is a neuropeptide involved in animal and human reproductive and social behavior, with implications for a range of psychiatric disorders. However, the therapeutic potential of oxytocin in mental health care suggested by animal research has not been successfully translated into clinical practice. This may be partly due to a poor understanding of the expression and distribution of the oxytocin signaling pathway in the human brain, and its complex interactions with other biological systems. Among the genes involved in the oxytocin signaling pathway, three genes have been frequently implicated in human social behavior: *OXT* (structural gene for oxytocin), *OXTR* (oxytocin receptor), and *CD38* (central oxytocin secretion). We characterized the distribution of *OXT, OXTR,* and *CD38* mRNA across the brain, identified putative gene pathway interactions by comparing gene expression patterns across 20737 genes, and assessed associations between gene expression patterns and mental states via large-scale fMRI metaanalysis. In line with the animal literature, expression of the three selected oxytocin pathway genes was increased in central, temporal, and olfactory regions. Across the brain, there was high co-expression with several dopaminergic and muscarinic acetylcholine genes, reflecting an anatomical basis for critical gene pathway interactions. Finally, fMRI meta-analysis revealed that the oxytocin pathway gene maps correspond with motivation and emotion processing.

## Introduction

Oxytocin is an evolutionarily conserved neuropeptide implicated in an array of social and reproductive behaviors, and its role in complex behavioral traits and in the pathophysiology of mental health conditions has attracted considerable attention (1). Oxytocin is mostly synthesized in neurons in the supraoptic nucleus and the paraventricular nucleus, and released both systemically and centrally. Human research has shown beneficial effects of intranasal oxytocin on performance on tests assessing social cognition (2), gaze to the eye region (3), and the retrieval of social cues (4). Moreover, single nucleotide polymorphisms in oxytocin pathway genes have been associated with social behavior and psychiatric disorders (5). Emerging evidence also points to the oxytocin system’s role in energy metabolism (6), with relevance for metabolic functions (7).

While central oxytocin receptor (*OXTR*) mRNA localization in the rodent brain is well-described (8), its anatomical distribution across the human brain is poorly understood, as investigations have tended to sample very few brain regions (9, 10). Partly attributed to this limited understanding of human central *OXTR* gene expression (11), the translational promise of oxytocin provided by preliminary clinical trials (e.g., 12, 13) has yet to be fulfilled. The distribution of *OXTR* mRNA across the brain provides a proxy for the distribution of central oxytocin binding (14, 15), allowing for a detailed mapping of the anatomical geography of the oxytocin system in the human brain. Seminal animal work using histochemistry and immunohistochemistry revealed high concentrations of *OXTR* mRNA in the hypothalamus, amygdala, olfactory bulb, ventral pallidum, and the dorsal vagal nucleus (16, 17). Further, experimentally increasing (18) or decreasing (19) *OXTR* expression in the prairie vole nucleus accumbens modulated partner preference behavior, suggesting an intriguing correspondence between the spatial distribution of *OXTR* mRNA, its functional neuroanatomy, and behavioral relevance.

In addition to the anatomical distribution of *OXTR* mRNA, characterizing its interactions with other key elements of the oxytocin signaling pathway and biological systems beyond this pathway is critical for determining oxytocin’s behavioral and functional relevance. Along with *OXTR, CD38* and oxytocin-neurophysin I (*OXT*) genes in the oxytocin signaling pathway have been implicated in human social behavior (5). Specifically, *CD38* is involved in central oxytocin secretion20 (20), and *OXT* encodes the oxytocin prepropeptide containing the nonapeptide oxytocin and the carrier protein neurophysin-I (21). *OXT* mRNA is highly expressed in human paraventricular nucleus of the hypothalamus, the lateral hypothalamic area, and the supraoptic nucleus, and there is evidence of co-expression with *OXTR* mRNA and the μ and κ types opioid receptor mRNA22 (22), providing a putative avenue for interactions between the oxytocin and opioid pathways.

The interactions with the oxytocin system extend beyond the opioid pathway, including the dopaminergic (23, 24) and muscarinic acetylcholine (9, 25) circuits, with possible implications for social behavior and psychiatric disorders. For instance, the dopamine D2-receptor subtype (DRD2), has been implicated in various putative intermediate phenotypes in psychiatric illness, including motivational processing (26) and pair bonding in animal models (27, 28). Moreover, the muscarinic acetylcholine M4 receptor (*CHRM4*) has been associated with schizophrenia (29) and implicated in cognitive flexibility (30), social impairment (31), and dopamine release (32). Finally, oxytocin can also bind to *AVPR1A* receptors (33), which has also been linked to social functioning (34). However, the mRNA co-expression of these systems, and potentially others, with the oxytocin system is not well characterized.

While brain regions with high oxytocin pathway gene expression in humans have been identified (9, 10, 22), inference of specific mental states from single brain regions is elusive. For instance, commonly observed increases in medial and lateral frontal region activity during emotion and pain processing seem to be better explained by more general sustained attention processes (35). Leveraging data from more than 11,000 fMRI studies, NeuroSynth (35) allows for reverse inference of mental states based on a given brain gene expression map with high specificity. Establishing the specific mental state correlates of oxytocin pathway genes will provide a deeper understanding of the central human oxytocin system and its relevance for brain functions and mental health.

By leveraging the Allen Human Brain Atlas, which offers a uniquely comprehensive gene expression survey from six neurotypical adult human brains, we first characterize the anatomical distribution of mRNA expression of *OXT, OXTR* and *CD38*, which are key constituents in the oxytocin pathway. Second, we explore putative gene interactions by identifying mRNA maps with overlapping anatomical distributions with our target genes across all protein coding genes in the database, additionally focusing on selected dopamine, muscarinic acetylcholine, and vasopressin gene sets. Third, we decode the mental state relevance of the selected oxytocin genes using quantitative reverse inference via large-scale fMRI meta-analysis, and assess specificity across 20737 mRNA maps.

## Materials and Methods

### Post-mortem brain samples

mRNA distribution data was collected from the Allen Human Brain Atlas (http://human.brain-map.org). Three donors were Caucasian males, one donor was a Hispanic female and two donors were African-American males. Mean donor age was 42.5 (S.D.=11.2) years. Data was collected 22.3 (S.D.=4.5) hours after demise, on average (See Table S1 for details). Each brain was sampled in 363-946 distinct locations, either in the left hemisphere only (*n* = 6), or over both hemispheres (*n* = 2) using a custom Agilent 8 x 60K cDNA array chip. Institutional review board approval was obtained at each tissue bank and repository that provided tissue samples, and informed consent was provided by each donor’s next-of-kin. For more details regarding procedures and data collection, see http://help.brain-map.org/display/humanbrain/Documentation.

### Data extraction

The full dataset of protein coding genes (*n* = 20737) were extracted from the Allen Human Brain Atlas. Of primary interest were the three oxytocin pathway genes that have been the predominant targets in previous studies of social behavior: *OXTR, CD38,* and *OXT* (5). Three other selected sets of mRNA, which are thought to co-express with central oxytocin pathway mRNA and modulate social behavior, were also of specific interest: A dopaminergic set (*DRD1, DRD2, DRD3, DRD4, DRD5, COMT, DAT1*) (23, 24, 36), a muscarinic acetylcholine set (*CRHM1, CRHM2, CRHM3, CRHM4,CRHM5*) (9, 25), and a vasopressin set *(AVPR1A, AVPR1B)* (34, 37). Of additional interest were complete sets of dopaminergic (*n* = 63), cholinergic (*n* = 79), and oxytocinergic (*n* = 94) genes, which were identified using the Kyoto Encyclopaedia of Genes and Genomes (KEGG) database (http://www.kegg.jp) (see Dataset S1 for a full list). If more than one probe for each mRNA was available, we selected the probe with the highest signal-to-noise ratio (mean/standard deviation), which represented the probe with least amount of spatial variability among donors (Dataset S2).

### mRNA expression maps

While the Allen Human Brain Atlas is a comprehensive dataset, expression data is missing from some regions or not available from all donor samples. To assess accurate cognitive correlates of gene expression using meta-analysis via the NeuroSynth tool, voxel-by-voxel mRNA maps provide optimal information. To create these voxel-by-voxel volumetric expression maps, we first marked all the sample locations and expression values in native image space. To interpolate missing voxels, we labeled brain borders with the sample expression value that had the closest distance to a given border point (Fig. S1a). Next, we linearly interpolated the space between scattered points by dividing that space into simplices based on Delaunay triangulation (Fig. S1b), then linearly interpolated each simplex with values to yield a completed map (Fig. S1c). All maps were computed in Matlab 2014a (The Mathworks Inc., Natick, MA, USA). Next, we registered all the individual brains to the MNI152 (Montreal Neurological Institute) template using ANTs (Advanced Normalization Tools; 38). To extract region specific statistics for 54 brain regions, we used regions extracted based on the Automated Anatomical Label (AAL) atlas.

For out-of-sample validation, central oxytocin pathway median expression profiles for 10 distinct brain regions were extracted from the Genotype-Tissue Expression (GTEx) project database (39). Although this dataset offers less spatial precision, the data is derived from a larger dataset of donors (mean sample size for mRNA expression across brain regions = 131.7, range = 88 to 173). For this comparison, median gene expression for these 10 distinct GTEx regions were calculated using Allen dataset (Fig. S2). Spearman’s rank correlation coefficients *(r_s_)* were calculated to assess the relationship between oxytocin pathway expression profiles from the two databases. Sex differences in central oxytocin pathway gene expression were also examined in the GTEx sample.

### Central gene expression

Statistical analyses were performed with R statistical package (version 3.3.2). Across the brain, gene expression for the right hemisphere samples was highly correlated with the corresponding left hemisphere samples for gene expression values for oxytocin pathway genes (*OXTR: r_s_* = 0.89, *p* < 0.001 *CD38: r_s_* = 0.87, *p* < 0.001; *OXT: r_s_* = 0.78, *p* < 0.001). Thus, the left hemisphere was used for analysis given its higher sample size. One-sample *t*-tests (two-tailed) were conducted to assess which of the 54 left hemisphere regions from six donor samples expressed mRNA to a significantly greater or lesser degree compared to average mRNA expression across the brain. To correct for multiple tests (54 in total), reported p-values were adjusted using a false discovery rate (FDR) threshold. Cohen’s *d* values for one-sample *t*-tests were calculated to yield a measure of effect size.

Based on raw expression data, we generated a 17 x 17 correlation matrix for each donor including the following selected genes (Oxytocin pathway set: *OXTR, CD38, OXT;* Dopaminergic set: *DRD1, DRD2, DRD3, DRD4, DRD5, COMT, DAT1;* muscarinic acetylcholine set: *CRHM1, CRHM2, CRHM3, CRHM4, CRHM5;* vasopressin set: *AVPR1A, AVPR1B)* and then averaged these matrices over the six donors. This mean correlation matrix was clustered using the complete linkage method (40) to assess coexpression patterns. To assess putative gene-gene interactions beyond selected gene sets, we computed the spatial Spearman’s correlations between all available protein coding mRNA probes (n = 20737) and three oxytocin pathway mRNA probes. Finally, to explore whether these coexpression patterns are not driven by global expression differences between cortical and non-cortical areas, we visualized the similarities in expression patterns based on cortical areas only.

### Mental state term correlates

To identify the mental state correlates of mRNA brain expression maps, we performed quantitative reverse inference via large-scale meta-analysis of functional neuroimaging data using the NeuroSynth framework (35). We created a composite brain map representing an average of 6 individuals on the left hemisphere. Using the NeuroSynth framework, we correlated our mRNA maps with specificity Z maps, reflecting the posterior probability maps, which display the likelihood of a given mental state term being used in a study if activation is observed at a particular voxel.

This analysis was based on text-mining a corpus of 11,406 fMRI studies. We assessed the 15 strongest relationships (Spearman’s *r*) between *OXTR, CD38,* and *OXT* expression maps (5 unique terms for each gene) and all available mental state (i.e., non-anatomical) maps available in the NeuroSynth database. To test for specificity, we also assessed the relationship between these 15 mental state maps and the 20737 mRNA maps. We plotted these distributions and identified the rank of *OXTR, CD38,* and *OXT* Spearman’s *r* value compared to all other associations.

### Data and analysis script availability

mRNA expression data is available from Allen Human Brain Atlas (http://human.brain-map.org) and GTEx (http://gtexportal.org). The Matlab script for producing the brain region-specific data, the resulting dataset, and the R script used for statistical analysis is available at href="https://osf.io/jp6zs/.

## Results

### Central gene expression

Compared to average expression across the brain, high expression of *OXTR* mRNA levels was observed in the olfactory bulbs (*p* = .001, *d* = 7.3), caudate (*p* = .001, *d* = 5.3), putamen (*p* = .003, *d* = 3.6), pallidum (*p* = .007, *d* = 2.9), amygdala (*p* = .01, *d* = 2.4), parahippocampal region (*p* = .03, *d* = 1.9), and hippocampus (*p* =.03, *d* = 1.8). There was lower *OXTR* expression in the cerebral crus I (*p* = .02, *d* = 2.1), cerebral crus II (*p* = .003, *d* = 3.7), cerebellum 7b (*p* = .003, *d* = 3.6), and cerebellum 8 (*p* = .002, *d* = 4.6; Fig. 1a; Dataset S3). Central *OXTR* expression patterns were comparable to patterns in the GTEx database *(r_s_* = 0.77, *p* = 0.01; Fig. S3). *CD38* gene expression was higher than average in the caudate (*p* = .003, *d* = 4.4), pallidum (*p* = .003, *d* = 4.4) olfactory bulbs (*p* = .01, *d* = 3.1), putamen (*p* = .02, *d* = 2.8), thalamus (*p* = .04, *d* = 2.1), and cingulate anterior (*p* = .047, *d* = 1.8; Fig. 1b). There was lower *CD38* expression in the middle temporal pole (*p* = .047, *d* = 1.9) and cerebellum 7b (*p* = .047, *d* = 1.8). Central *CD38* expression was comparable to *CD38* central expression in the GTEx database (r_s_= 0.68, *p* = 0.04). High expression of *OXT* mRNA was observed in the pallidum (*p* = .03, *d* = 3.3), and the thalamus (*p* = .03, *d* = 2.7; Fig. S4). The rank order of central *OXT* expression was related to the GTEx database rank order, but this fell short of statistical significance *(r_s_* = 0. 42, *p* = 0.23). Expression of genes included in the selected dopaminergic, muscarinic acetylcholine, and vasopressin sets are presented in Figures S5, S6, S7, respectively, and Dataset S3. Of note, *DRD2, COMT, and CHRM4* genes were highly expressed in central regions. Inspection of gene expression data from the GTEx database suggested no sex differences in central oxytocin pathway gene expression (Fig. S8).

**Figure 1.**
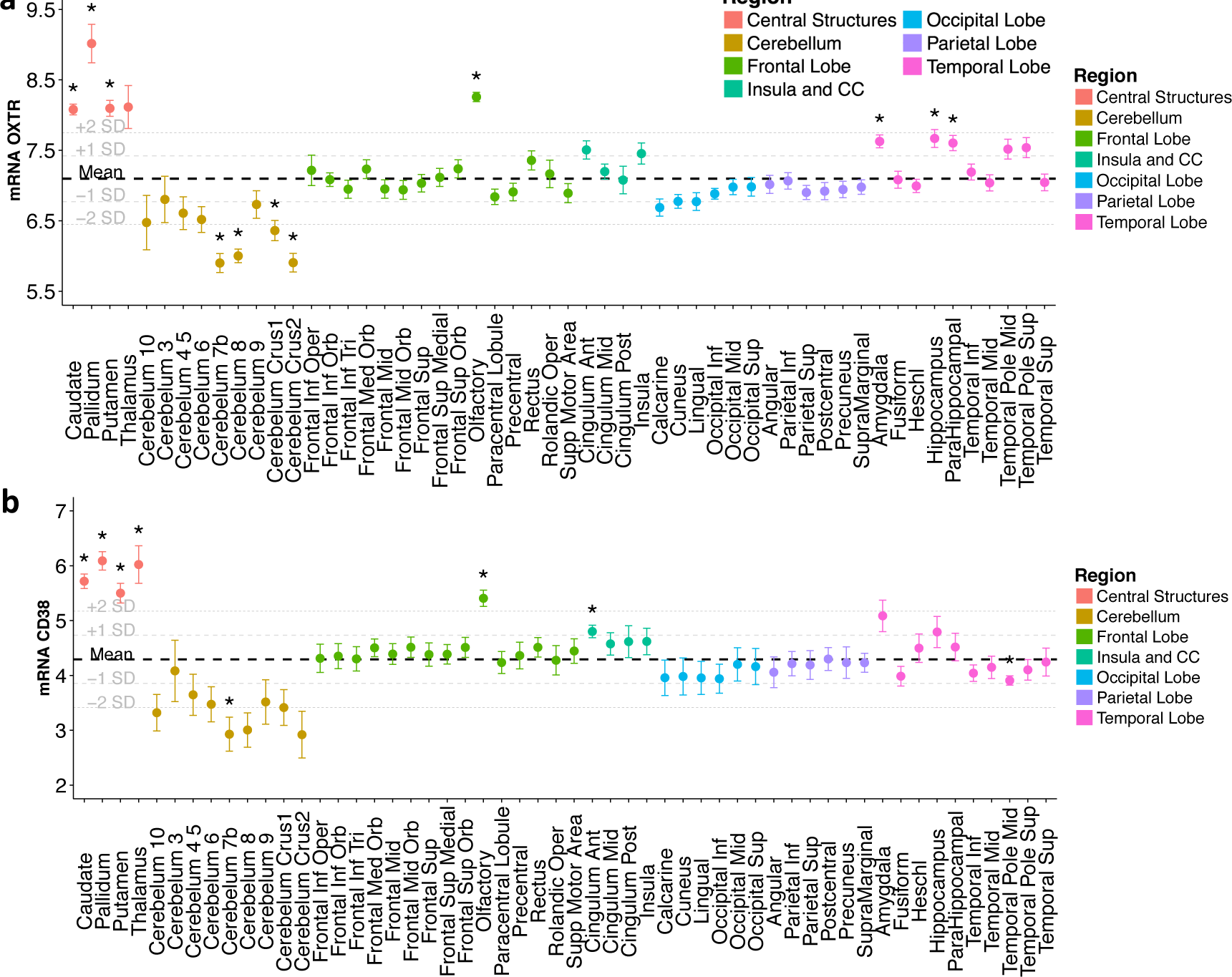
Central gene expression for *OXTR* (a) and *CD38* (b). Each point represents mean expression with standard errors for a given brain region. Data was collected from the Allen Human Brain Atlas, representing mean expression across six donors, and then voxel-by-voxel volumetric expression maps were created (Fig. S1). Each brain was registered the Montreal Neurological Institute template using Advanced Normalization tools. Statistics for each region were extracted using based on the Automated Anatomical Label atlas. The bolded dashed lines represent mean expression across the all regions with 1 and 2 standard deviations also shown. One sample t-tests were performed to assess which regions expressed mRNA to a significantly greater degree compared to average expression across the brain. Analyses suggest that compared to average brain expression, there is increased expression of *OXTR* and *CD38* in central and temporal brain structures, along with the olfactory region. Lower than average expression is observed in the cerebellum. Central *OXT* expression patterns are shown in Figure S4. *p < 0.05 (FDR corrected for 54 tests).

A Spearman’s correlation matrix demonstrating the relationships between the genes of interest using the raw expression data is presented in Figure 2a. Hierarchical clustering revealed that that the oxytocin pathway genes (*OXTR, CD38, OXT*) cluster with elements of the dopaminergic system (*DRD1, DAT1, COMT, DRD2*), *CHRM4* and *CHRM5*. The correlation of each of these genes within its complete gene set family is visualized in Figure 2b. A comparison of expression patterns across the all brain regions and only cortical samples suggest that *OXTR* and *CD38* relationships appear to be global, whereas *OXT* co-expression seems to be driven by correlations within cortical areas (Fig. S9). Histograms of the correlations between the three oxytocin pathway mRNA probes and all other available mRNA probes are presented in Figure 3. Exploratory correlation analyses with all available 20737 protein coding gene probes in the Allen Human Brain Atlas and each of the three oxytocin pathway gene probes are summarized in Table S2, with the top 10 strongest positive and top 10 strongest negative correlations. The top positive mRNA map correlations for *OXTR* were *NTSR2 (r_s_* = 0.8), *GLUD2* (*r_s_* = 0.78), and *GLUD1* (*r_s_* = 0.78).

**Figure 2.**
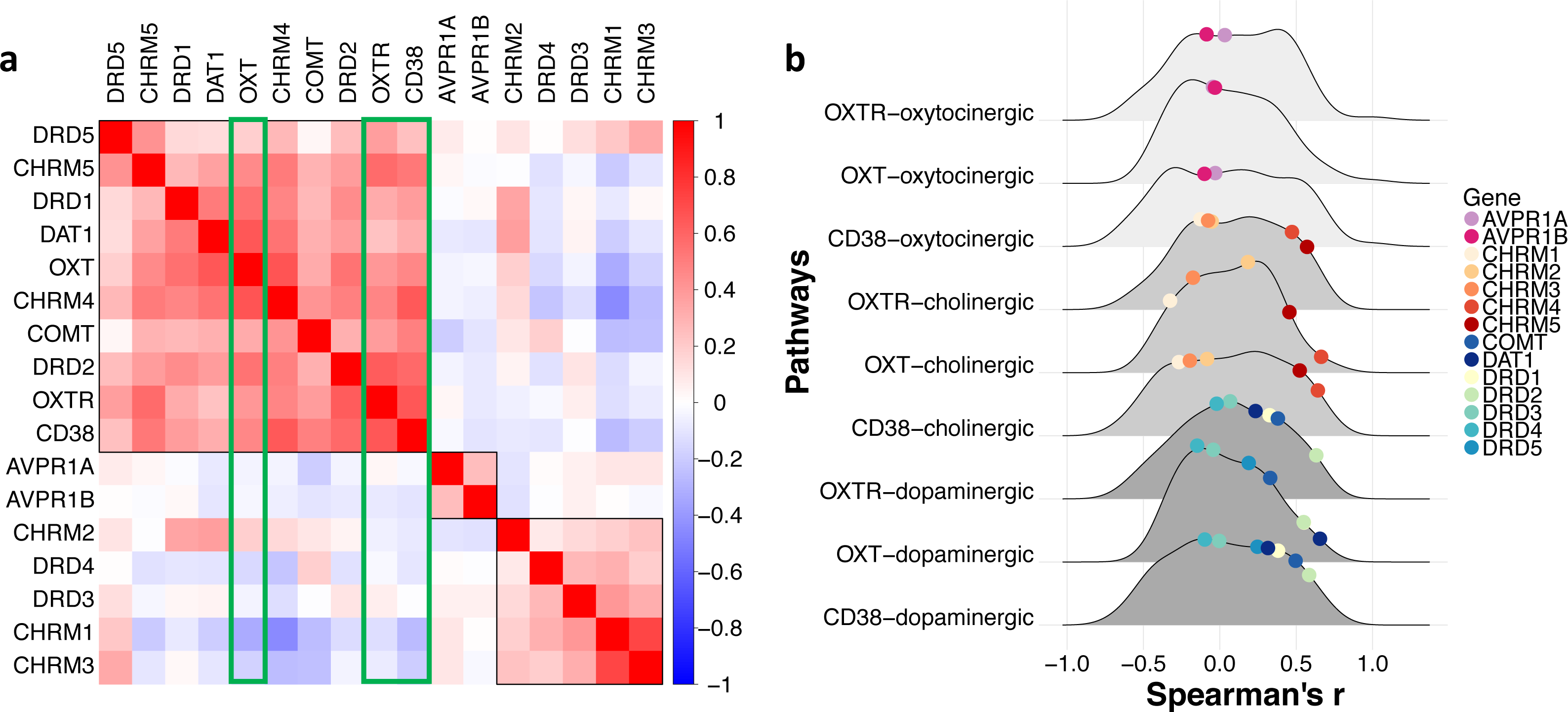
Central co-expression of selected oxytocin, dopaminergic, muscarinic acetylcholine, and vasopressin gene sets. (a) Co-expression patterning for the central expression of selected oxytocin, dopaminergic, muscarinic acetylcholine, and vasopressin gene sets. The complete linkage method was used to identify 3 clustering groups (black squares). The three oxytocin pathway genes (green rectangles) are clustered together, along with *DRD2, COMT, CHRM4, DAT1, DRD1, CHRM5,* and *DRD5* genes. (b) Central co-expression of *OXTR, CD38,* and *OXT* with selected vasopressin, muscarinic acetylcholine, and dopaminergic gene sets. Spearman’s correlations are visualized on a density distribution demonstrating all coexpression correlations between the of *OXTR, CD38,* and *OXT* and all genes in the complete oxytocinergic (n = 94), cholinergic (n = 79), and dopaminergic (n = 63) gene sets (see Dataset S1 for full lists).

**Figure 3.**
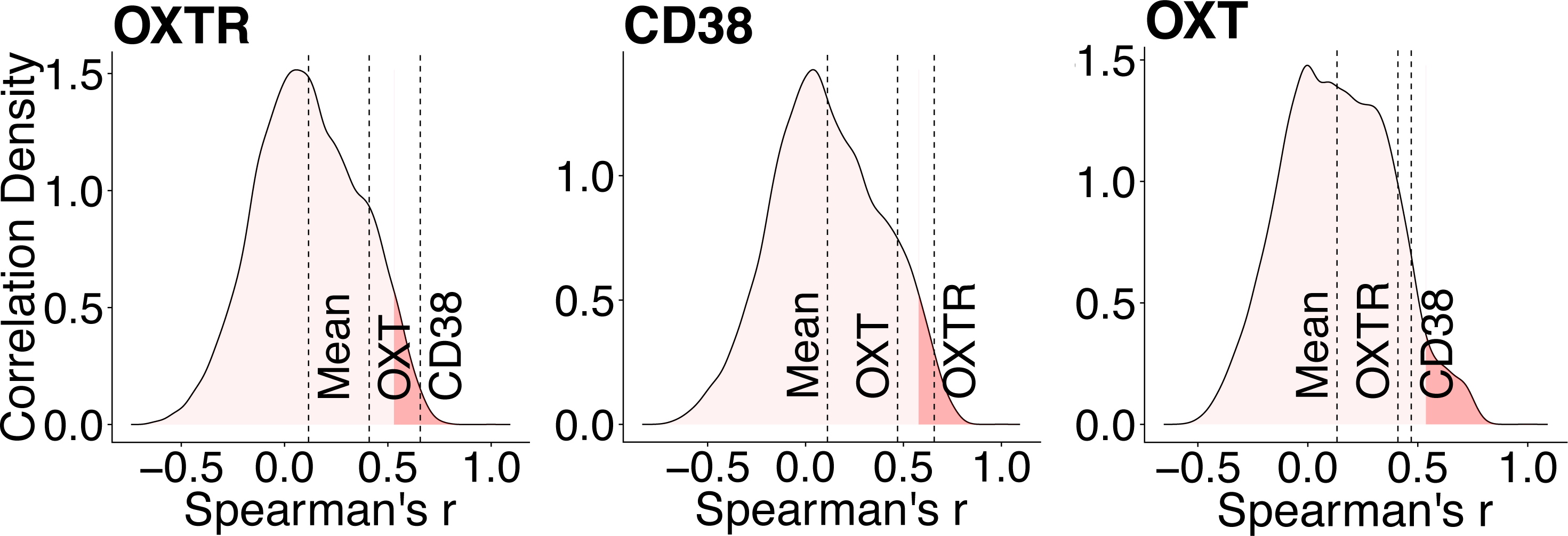
Co-expression of oxytocin pathway genes against 20737 protein coding genes. Density plots visualize the distributions of correlations between *OXTR, CD38,* and *OXT,* and all other mRNA probes. For each oxytocin gene, the position of the other two oxytocin genes is shown. Dark red portions of the distributions represent the top 5% of correlations.

### Cognitive correlates

Decoding mental states meta-analytically from mRNA maps (Fig. 4) via quantitative reverse inference revealed that *OXTR, CD38, and OXT* mRNA expression maps were most highly correlated with functional imaging maps with motivation-related topics (e.g., “reward”, “anticipation”, “incentive”), emotion-processing, and aversive-related topics (Fig. 5a, Table S3). Figure 5b shows the full distribution of correlation coefficients for each mental state term across all 20737 gene maps, with labelled oxytocin pathway genes and their absolute rank for each term (Table S3). Notably, the *OXTR* map was ranked among the top 100 out of 20737 genes for several functional imaging maps (i.e., the top 0.5%).

**Figure 4.**
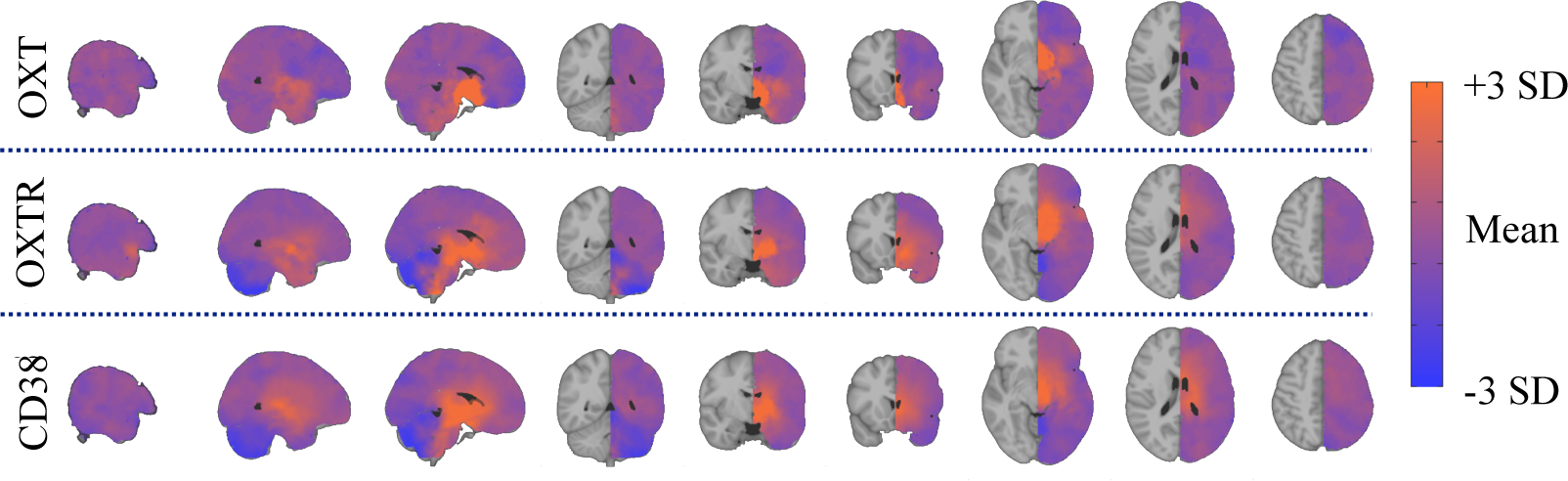
Brain gene expression maps for *OXT*, *OXTR*, and *CD38*. Voxel-by-voxel volumetric mRNA expression maps were created using left hemisphere data from the Allen Human Brain Atlas dataset. SD = standard deviation.

**Figure 5.**
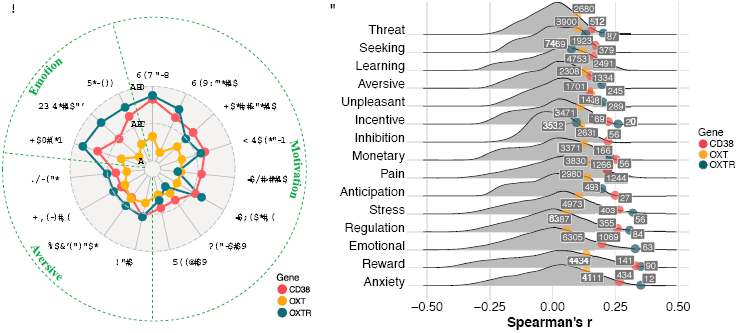
Cognitive correlates of central oxytocin pathway gene expression. Mental states were meta-analytically decoded from central oxytocin pathway mRNA maps using the NeuroSynth framework. (a) The top five strongest relationships for *OXTR, CD38,* and *OXT* (Spearman’s r) are shown, with duplicates removed. (b) The distribution of Spearman correlations between each protein coding gene map (*n* = 20737) and mental state maps. The absolute ranking for each oxytocin pathway gene out of 20737 correlations are shown (also see Table S3).

## Discussion

The anatomical distribution of gene expression in the brain is heterogeneous and highly coordinated (41). Whereas dynamic alterations in gene expression are essential in response to environmental demands, and are critically involved in a range of cognitive functions, learning, and diseases (42, 43), the basic organization likely partly reflects the evolutionary conserved modular layout of the brain (44). Indeed, gene-gene co-expression patterns form specific genetic signatures in the brain, representing distinct tissues and biological systems (41), and likely reflect the potential of complex and differential gene-gene interactions with implications for brain disorders and mental health. Here, we leveraged the unique human brain mRNA expression library from Allen Brain Atlas to show that mRNA reflecting specific genes in the oxytocin pathway are highly expressed in central and temporal brain structures, along with the olfactory region. We also show reduced expression of *OXTR* and *CD38* in the cerebellum, consistent with prior animal research (45). Importantly, the observed oxytocin pathway expression patterns from the Allen database, particularly OXTR and CD38, were consistent with oxytocin pathway expression patterns observed in the GTEx database.

Oxytocin pathway genes showed considerable co-expression with *DRD2, COMT, DAT1,* and *CHRM4* genes, providing evidence for putative interactions between dopaminergic and muscarinic acetylcholine systems with the central oxytocin pathways. Exploratory analysis between oxytocin pathway mRNA and 20737 mRNA probes revealed several relationships worth noting in the context of metabolic and feeding regulation, as well as psychiatric disorders. We discovered that *OXTR* is highly co-expressed with Neurotensin receptor gene *(NTSR2),* which has been found to regulate ethanol consumption in mice (46), and Glutamate dehydrogenase 1 and 2 *(GLUD1, GLUD2),* which are involved in energy homeostasis and insulin secretion (47-49). Second, *CD38* is highly co-expressed with *NTSR2, GLUD1,* and *SPX* (Spexin), with the latter associated with weight regulation (50).

These results are consistent with emerging evidence that the oxytocin system may play a role in the metabolic and feeding dysregulation often observed in severe mental illnesses (7). In regards to psychiatric disorders, there were also strong negative correlations between oxytocin pathway genes and *CADPS2,* which has been associated with autism (51) and autistic-like phenotypes in mice (52), *PAK7* and *RTN4R* (Nogo-66), which have been associated with schizophrenia (53, 54), and *GALNT17* (WBSCR17), which has been associated with William’s syndrome (55). Quantitative reverse inference via meta-analysis of 11,406 fMRI studies also revealed that the distribution of the central genetic oxytocin pathway is most strongly associated with terms related to motivation and emotion, supporting the gene-system interactions suggested by the gene co-expression patterns.

Compelling rodent evidence suggests that direct central oxytocin administration influences social behavior and cognition via action on central oxytocin receptors (56). In these animals, oxytocin receptors are located in regions that are crucial for social behavior and the processing of social cues (8). Our data reveals striking similarities with central oxytocin pathway mRNA distribution observed in rodents (16, 17) and non-human primates (57). Additionally, our data provides evidence that oxytocin may exert its effects synergistically with the dopaminergic network (23). Indeed, the oxytocin and dopaminergic systems work in concert to promote rodent maternal behaviors (58) and central D2 receptors modulate social functioning in prairie voles (59, 60). There is also evidence of direct dopaminergic fibers to the supraoptic nucleus and the paraventricular nucleus (61), consistent with demonstrations that dopamine may stimulate oxytocin release in the hypothalamus (62). This is also congruous with reported deficits in dopaminergic signaling in schizophrenia (63) and autism (64). The muscarinic acetylcholine system may also work with the oxytocin system to boost attention (65) and to assist with the release of oxytocin (66). Relatedly, the muscarinic acetylcholine system is thought to contribute to the cognitive dysfunction observed in schizophrenia (67). Altogether, these findings are consistent with a role of oxytocin in the processing of social stimuli and corroborate emerging evidence for interactions between the oxytocin, acetylcholine (9), and dopaminergic systems (23), and for oxytocin’s role in metabolic regulation (7).

Our data revealed high expression of oxytocin pathway mRNA in the olfactory region, which is in line with animal research (e.g., 68). Indeed, social olfactory deficits in mice without the oxytocin gene (69) are rescued with injection of oxytocin into rat olfactory bulbs (70). Increased mRNA in the human olfactory region may be vestigial, as olfaction is not as important for human conspecific identification compared to most other mammals due to species specialization, however, intranasal oxytocin has been shown to improve olfactory detection performance in schizophrenia (71), and intranasal oxytocin is thought to enter the brain via the olfactory bulbs (72).

There are some important limitations worth noting. First, given the nature of human tissue collection for mRNA studies, the sample size was small. However, the relatively small amount of measurement error for the mRNA probes indicates high precision. Second, there was only 1 female in the sample, which prohibited an examination of sex differences in mRNA distribution from the Allen dataset. However, data from the GTEx dataset demonstrated little evidence of sex differences in central oxytocin pathway gene expression, consistent with preliminary human research (9). Third, we analyzed data derived from dissected tissue that contained multiple cell types, thus diluting the transcriptional contribution and dynamic range of expression of any one specific cell type. *OXT* mRNA has been identified in hypothalamic axons (73), however, current limitations prevent cell-type-specific approaches for systematically analyzing the central transcriptome. Relatedly, it is important to note that the relationship between mRNA and protein levels is not always linear nor translated into apparent phenotypic differences. Moreover, as our results only describe an association between gene transcript locations and voxel-based neural activity patterns that correspond to various mental states, it is possible that other neuronal types underlie this neural activity. Finally, despite animal models demonstrating that *OXTR* expression levels directly modulate social behavior (18, 19), it remains unclear whether the increased expression of *OXTR* as measured by the presence of its transcript in our human sample is directly relevant for the activation of the receptor during the relevant mental state.

It has been suggested that comprehensive clinical trials need to demonstrate engagement of drug targets, such as *OXTR* occupancy reflected by regional brain activity changes (11). Without precise targets, it is unclear whether non-significant effects of intranasal oxytocin, beyond insufficient statistical power (74), are due to an inefficacious drug or misidentified drug targets. By identifying accurate oxytocin pathway targets in the human brain, our study provides a tentative oxytocin treatment target map that may help facilitate efforts to better understand oxytocin treatment efficacy (11, 72). Analysis of data from the Allen Brain Atlas database provides a unique opportunity to map the central oxytocin system and explore co-expression with other systems. Altogether, these results provide a proof-of-principle demonstration of corresponding cognitive and gene expression patterns of a neuropeptide pathway involved in complex human behaviors.

## Acknowledgements

This research was supported by an Excellence Grant from the Novo Nordisk Foundation to DSQ (NNF16OC0019856), Research Council of Norway grants to OAA (223273) and LTW (249795), a KG Jebsen Stiftelsen grant to OAA (SKGJ-MED-008), and South-Eastern Norway Regional Health Authority grants to OAA (2017-112) and LTW (2014097, 2015073).

